# *De novo* genome assembly of the land snail *Candidula unifasciata* (Mollusca: Gastropoda)

**DOI:** 10.1101/2021.01.23.427926

**Authors:** Luis J. Chueca, Tilman Schell, Markus Pfenninger

## Abstract

Among all molluscs, land snails are an economically and scientifically interesting group comprising edible species, alien species and agricultural pests. Yet, despite its high diversity, the number of whole genomes publicly available is still scarce. Here, we present the draft genome assembly of the land snail *Candidula unifasciata*, a widely distributed species along central Europe, which belongs to Geomitridae family, a group highly diversified in the Western-Palearctic region. We performed a whole genome sequencing, assembly and annotation of an adult specimen based on PacBio and Oxford Nanopore long read sequences as well as Illumina data. A genome of about 1.29 Gb was generated with a N50 length of 246 kb. More than 60% of the assembled genome was identified as repetitive elements, and 22,464 protein-coding genes were identified in the genome, where the 62.27% were functionally annotated. This is the first assembled and annotated genome for a geometrid snail and will serve as reference for further evolutionary, genomic and population genetic studies of this important and interesting group.

## 1. Introduction

Gastropods are the largest group among molluscs, representing almost the 80% of the species. Although most of the them are present in marine habitats, land snails diversity is estimated around 35.000 species (Solem 1984). Due to its low dispersal abilities, land snails have been employed in many evolutionary and population genomics studies (Stankowski 2013; Schilthuizen and Kellermann 2014; Chueca *et al*. 2017; Haponski *et al*. 2017). While these studies are mainly based on few loci, transcriptomes or mitochondrial genomes (Kang *et al*. 2016; Romero *et al*. 2016; Razkin *et al*. 2016; Korábek *et al*. 2019), only a couple of whole nuclear genomes of land snails species are available so far. Geomitridae is one of the most diverse families of molluscs in Western-Palearctic region. The family is composed by small to medium-size species, characterized by presenting several reproductive adaptations to xeric habitats (Giusti and Manganelli 1987). *Candidula unifasciata* (NCBI:txid100452) is a land snail species widely distributed along western Europe, from southern France and Italy to central and northern Europe (Fig. 1). *C. unifasciata* inhabits dry meadows and open lowlands with rocks, being also present in gardens and vineyards. A recent molecular revision of *Candidula* (Chueca *et al*. 2018) revealed the polyphyly of the genus, and split the species that composed it into six genera, questioning the traditional anatomical classification. Although, there are many taxonomical, phylogeographical and evolutionary studies concerning Geomitridae species (Pfenninger and Magnin 2001; Sauer and Hausdorf 2010; Brozzo *et al*. 2020), the lack of reference genomes makes it difficult to investigate deeper biological and evolutionary questions about geomitrids and other land snails species. Here, we present the annotated draft genome of *Candidula unifasciata* that will be a valuable resource for future genomic research of this important taxonomic group.

**Figure 1.**
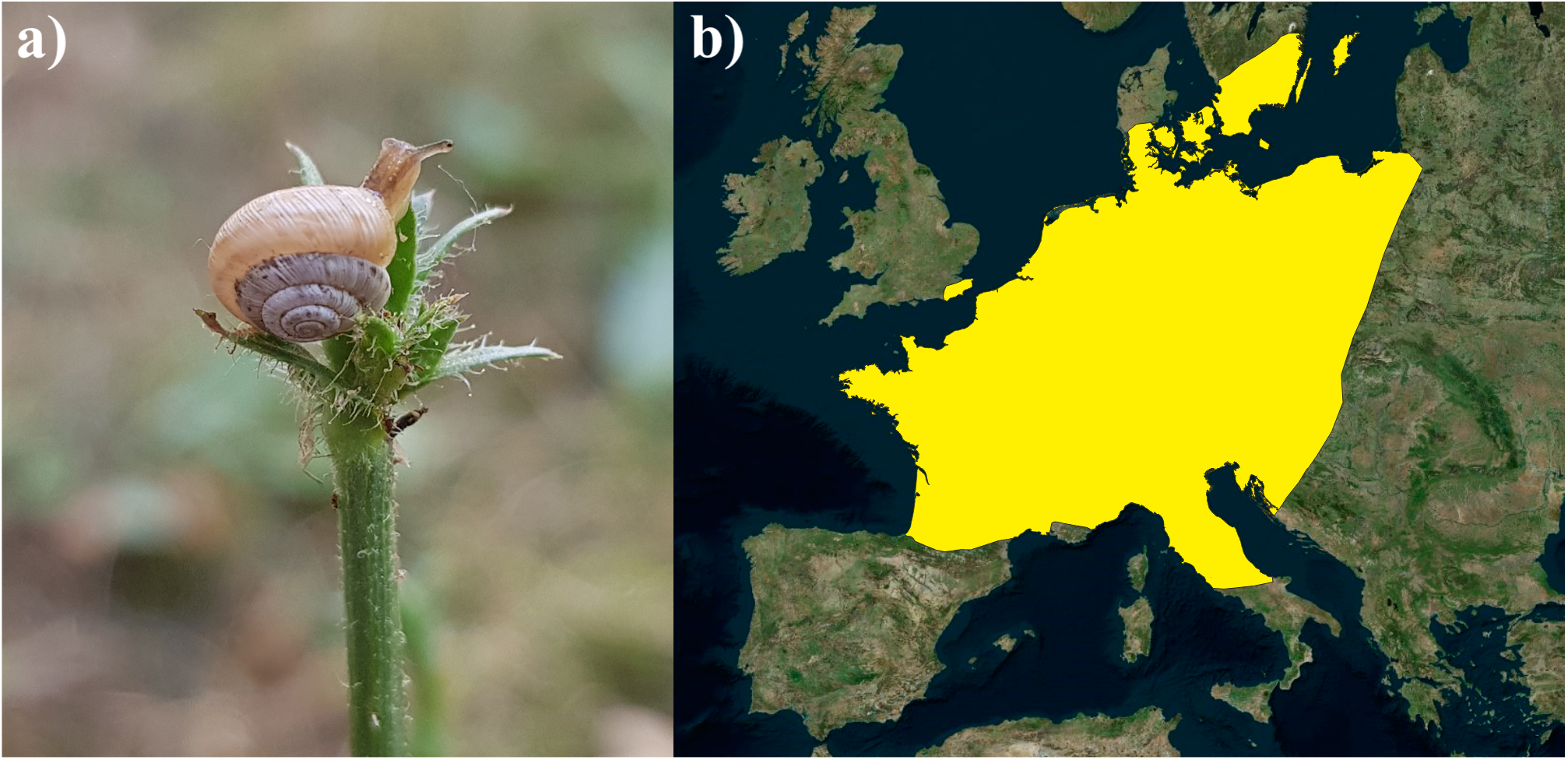
**a)** Picture of an adult specimen of *Candidula unifasciata*, copyright © Luis J. Chueca. **b)** Distribution range of *C. unifasciata* in Europe.

## 2. Materials and Methods

### 2.1 Sample collection, library construction, sequencing

A live population of *C. unifasciata* was collected from Winterscheid, Gilserberg, Gemany (50.93° N, 9.04° E). Genomic DNA was extracted from one specimen using the phenol/chloroform method and quality was checked by gel electrophoresis and NanoDrop ND-1000 spectrophotometer (LabTech, USA). A total of 5.6 μg of DNA was sent to Novogene (UK) for library preparation and sequencing. Then, a 300 base pair (bp) insert DNA libraries were generated using NEBNext® DNA Library Prep Kit and sequenced on 3 lanes of Illumina NovaSeq 6000 platform (150 bp paired-end [PE] reads). Quality of raw Illumina sequences was checked with FastQC (Andrews 2010). Low quality bases and adapter sequences were subsequently trimmed by Trimmomatic v0.39 (Bolger *et al*. 2014). For PacBio sequencing, a DNA library was prepared from 5 μg of DNA using the SMRTbell template prep kit v.1.0. Sequencing was carried out on 10 single-molecule real-time sequencing (SMRT) cells on an RSI instrument using P6-C4 chemistry.

To obtain Oxford Nanopore Technologies (ONT) long reads, we ran two flow cells on a MinION portable sequencer. Total genomic DNA was used for library preparation with the Ligation Sequencing kit (SQK-LSK109) from ONT, using the manufacturer’s protocols. Base calling of the reads from the two MinION flow cells was performed with guppy v4.0.11 (https://nanoporetech.com/nanopore-sequencing-data-analysis), under default settings. Afterwards, ONT reads quality was checked with Nanoplot v1.28.1 (https://github.com/wdecoster/NanoPlot) and reads shorter than 1000 bases and mean quality below seven were discarded by running Nanofilt v2.6.0 (https://github.com/wdecoster/nanofilt).

Two specimens, one adult and one juvenile, were ground together into small pieces using steel balls and a Retsch Mill. Then, RNA was extracted following an standard Trizol extraction. The integrity of total RNA extracted was assessed on an Agilent 4200 TapeStation (Agilent, USA), after which, approximately 1 µg of the total RNA was processed using the Universal Plus mRNA-seq library preparation kit (NuGEN, Redwood City, CA). Finally, the 300-bp insert size library was sequenced on a Illumina NovaSeq 6000 platform.

### 2.2 Genome size estimation

Genome size was estimated following a flow cytometry protocol with propidium iodide-stained nuclei described in (Hare and Johnston 2012). Foot tissue of one fresh adult sample of *C. unifasciata* and neural tissue of the internal reference standard *Acheta domesticus* (female, 1C = 2 Gb) was mixed and chopped with a razor blade in a petri dish containing 2 ml of ice-cold Galbraith buffer. The suspension was filtered through a 42-μm nylon mesh and stained with the intercalating fluorochrome propidium iodide (PI, Thermo Fisher Scientific) and treated with RNase II A (Sigma-Aldrich), each with a final concentration of 25 μg/ml. The mean red PI fluorescence signal of stained nuclei was quantified using a Beckman-Coulter CytoFLEX flow cytometer with a solid-state laser emitting at 488 nm. Fluorescence intensities of 5000 nuclei per sample were recorded.

We used the software CytExpert 2.3 for histogram analyses. The total quantity of DNA in the sample was calculated as the ratio of the mean red fluorescence signal of the 2C peak of the stained nuclei of the *C. unifasciata* sample divided by the mean fluorescence signal of the 2C peak of the reference standard times the 1C amount of DNA in the standard reference. Four replicates were measured to minimize possible random instrumental errors. Furthermore, we estimated the genome size by coverage from mapping reads used for genome assembly back to the assembly itself using backmap v0.3 (https://github.com/schellt/backmap; Schell *et al*. 2017). In brief, the method divides the number of mapped nucleotides by the mode of the coverage distribution. By doing so, the length of collapsed regions with many fold increased coverage is taken into account.

### 2.3 Genome assembly workflow

Different *de novo* genome assemblies were tested under different methods (see Table S1). The pipeline, which showed the best genome, was selected to continue further analyses. The draft genome was constructed from PacBio long reads using wtdbg2 v2.5 (Ruan and Li 2020), followed by three polishing rounds of Racon 1.4.3 (Vaser *et al*. 2017) and three polishing rounds of Pilon 1.23 (Walker *et al*. 2014). After that, Illumina and PacBio reads were aligned to the assembly using backmap.pl v0.3 to evaluate coverage distribution. Then, Purge Haplotigs (Roach *et al*. 2018) was employed, under default parameters and cut off values of 15, 72 and 160 to identify and remove redundant contigs.

### 2.4 Scaffolding and gap closing

To further improve the assembly, we applied two rounds of scaffolding and gap closing to the selected genome assembly. The genome was first scaffolded with the SMRT and ONT reads by LINKS v1.8.7 (Warren *et al*. 2015) and then with RNA reads by Rascaf v1.0.2 (Song *et al*. 2016). Long-Read Gapcloser v1.0 (Xu *et al*. 2018) was run three times after each scaffolding step, followed by three polishing rounds of Racon v1.4.3. BlobTools v.1.0 (Kumar *et al*. 2013; Laetsch and Blaxter 2017) was employed to screen genome assembly for potential contamination by evaluating coverage, GC content and sequence similarity against the NCBI nt database of each sequence. The resulting assembly was compared in terms of contiguity using Quast v5.0.2 (Gurevich *et al*. 2013), and evaluated for completeness by BUSCO v3.02 (Simão *et al*. 2015) against metazoa_odb9 data set.

### 2.5 Transcriptome assembly

RNA reads were also checked for quality and trimmed, as was explained above, and the transcriptome was assembled using Trinity v2.9.1 (Haas *et al*. 2013). Then, the transcriptome assembly was evaluated for completeness by BUSCO v3.0.2 against the against metazoa_odb9 data set. Moreover, the clean RNA-seq reads from different specimens were aligned against the reference genome by HISAT2 (Kim *et al*. 2015).

### 2.6 Repeat Annotation

RepeatModeler v2.0 (Smit and Hubley 2008) was run to construct a *de novo* repetitive library from the assembly. The resulting repetitive library created was employed by RepeatMasker v4.1.0 (http://www.repeatmasker.org/) to annotate and masked the genome.

### 2.7 Gene prediction and functional annotation

Genes were predicted by using different methods. First, genes models were predicted *ab initio* based on SNAP v. 2006-07-28 (Korf 2004) and the candidates coding regions within the assembled transcript were identified with TransDecoder v5.5.0 (https://github.com/TransDecoder/). Secondly, we used homology-based gene predictions by aligning protein sequences from SwissProt (2020-04) to the *Candidula unifasciata* masked genome with EXONERATE 2.2.0 (Slater and Birney 2005) and by running GeMoMa v1.7.1 (Keilwagen *et al*. 2016, 2018) taking five gastropods species as reference organisms. The selected species were *Pomacea canaliculata* (GCF_003073045.1; (Liu *et al*. 2018), *Aplysia californica* (GCF_000002075.1), *Elysia chlorotica* (GCA_003991915.1; (Cai *et al*. 2019), *Radix auricularia* (GCA_002072015.1; (Schell *et al*. 2017) and *Chrysomallon squamiferum* (GCA_012295275.1; (Sun *et al*. 2020), which were downloaded from NCBI. First, from the mapped RNA-seq reads, introns were extracted and filtered by the GeMoMa modules ERE and DenoiseIntrons. Then, we ran independently the module GeMoMa pipeline for each reference species using mmseqs2 and including the RNA-seq data. The five gene annotations were then combined into a final annotation file by using the GeMoMa modules GAF and AnnotationFinalizer. Finally, we aligned *C. unifasciata* transcripts against the masked genome using PASA v2.4.1 (Campbell *et al*. 2006) as implemented in autoAug.pl.

Gene prediction data from each method were combined using EVidenceMolder v1.1.1 (Haas *et al*. 2008) to obtain a consensus gene set for the raccoon-dog genome. Gene models from GeMoMa and SNAP were converted to EVM compatible gff3 files and combined with CDS identified by TransDecoder into a gene predictions file. After that, EVM was run including gene model predictions, protein and transcript alignments and repeat regions to produce a reliable consensus gene set.

Predicted genes were annotated by BLAST search against the Swiss-Prot database with an e-value cutoff of 10^−6^. InterProScan v5.39.77 (Quevillon *et al*. 2005) was used to predict motifs and domains, as well as Gene ontology (GO) terms.

## 3. Results and Discussion

### 3.1 Genome assembly

The calculated DNA content through flow cytometry experiments was 1.54 Gb. The genome size estimation by Illumina read coverage resulted in 1.42 Gb. The estimated heterozygosity by GenomeScope of the specimen employed for genome assembly was around 1.09% (Fig. 2.a), being in the range of other land snail genomes (Guo *et al*. 2019; Saenko *et al*. 2021). We generated sequence data for a total coverage of approximately 120.6X and 25.6X of Illumina and PacBio reads respectively. After scaffolding with long reads (PacBio and ONT) and RNA data, we produced a draft genome assembly of 1.29 Gb with 8,586 scaffolds and a scaffold N50 of 246 kb (Table 1). Completeness evaluation by BUSCO against the metazoan_odb9 data set showed high values, recovering more than the 92% as complete and less than the 6% as missing genes for both, assembly and annotation, analyses (Table 1). This results were in the range of other gastropods genome assemblies (Schell *et al*. 2017; Liu *et al*. 2018; Guo *et al*. 2019; Sun *et al*. 2020), being slightly better than closest relative assembly of *Cepaea nemoralis* (Saenko *et al*. 2021). For genome quality evaluation, we compared the *C. unifasciata* draft genome generated with other mollusc genomes publicly available. This comparison showed high quality in terms of contig number and scaffold N50 among land snail genomes. The mapping of the Illumina reads against the final genome assembly showed that the 98.56% of them were aligned to it, as well as a good removal of redundant contigs (Fig. 2b). Finally, BlobTools analysis didn’t reflect substantial contamination (Fig. 3), indicating the reliability of the data.

**Figure 2.**
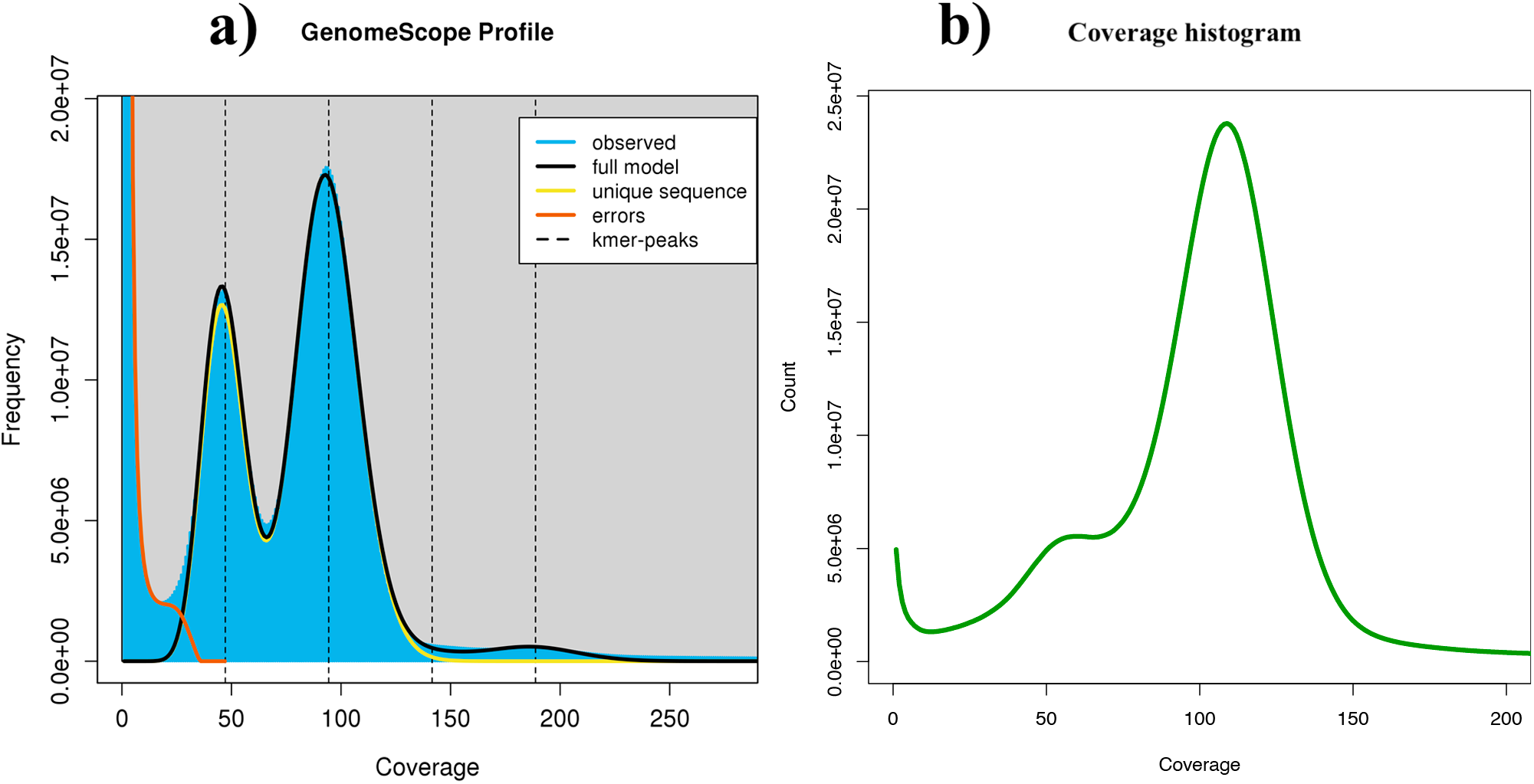
**a)** GenomeScope k-mer profile plot for *Candidula unifasciata* genome based on 21-mers in Illumina reads. **b)** Coverage histogram for the final assembly based on the Illumina reads.

**Table 1.** Genome assembly and annotation statistics for *C. unifasciata* and comparison with other land snails genomes.

| Statistic | *Candidula*<br>*unifasciata* | *Cepaea nemoralis* | *Achatina fulica* |
| --- | --- | --- | --- |
| Total sequence length | 1,286,461,591 | 3,490,924,950 | 1,850,322,141 |
| No. of contigs | 11,756 | 28,698 | 8,122 |
| Contig N50 | 246,413 | 330,079 | 721,038 |
| Contig L50 | 1,602 | 3,071 | 697 |
| No. of scaffolds | 8,586 | 28,537 | 921 |
| scaffolds > 10000 bp | 7,180 | 26,580 | 189 |
| Scaffold N50 | 246,413 | 333,110 | 59,589,303 |
| Scaffold L50 | 940 | 3,035 | 13 |
| GC content (%) | 40.69 | 41.25 | 38.77 |
| BUSCO complete (genome) | 92.4% (S:85.3%; D:7.1%) | 87.2% (S:74.3%; D:12.9%)* | 91.5% (S:84.6%; D:6.9%) |
| BUSCO fragmented (genome) | 1.6% | 3.8%* | 2.5% |
| BUSCO missing (genome) | 6.0% | 9.0%* | 6.0% |
| BUSCO complete (annotation) | 94.5% (S:86.0%; D:8.5%) | na | 95.6% (S:86.8%; D:8.8%) |
| **BUSCO fragmented (annotation)** | 2.6% | na | 1.9% |
| **BUSCO missing (annotation)** | 2.9% | na | 2.5% |
| **BUSCO complete (transcriptome)** | 94.7% (S:52.6%; D:42.1%) |  |  |
| **BUSCO fragmented (transcriptome)** | 3.8% |  |  |
| **BUSCO missing (transcriptome)** | 1.5% |  |  |

\*against metazoa\_odb10 dataset (n=954)

**Figure 3.**
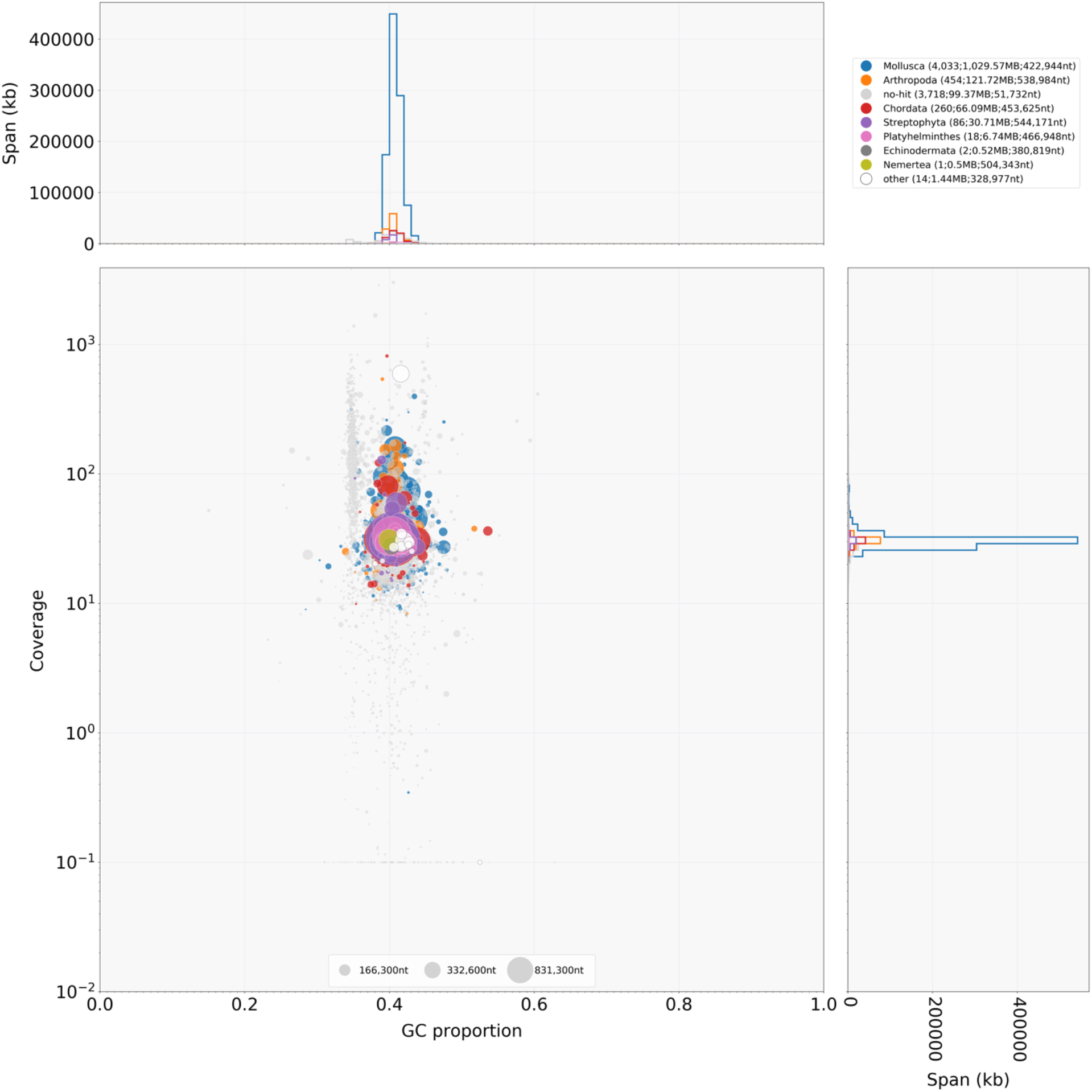
Blob plot showing read depth of coverage, GC content and size of each scaffold. Size of the blobs correspond to size of the scaffold and color corresponds to taxonomic assignment of BLAST.

### 3.2 Genome annotation

We estimated the total repeat content of the *C. unifasciata* genome assembly around 61.10% (Table 2), values slightly smaller than other land snails genomes (Guo *et al*. 2019; Saenko *et al*. 2021).

**Table 2.** Repeat statistics. *De novo* and homology based repeat annotations as reported by RepeatMasker and RepeatModeler for *C. unifasciata* and comparison with *Cepaea nemoralis*. Families of repeats included here are long interspersed nuclear elements (LINEs), short interspersed nuclear elements (SINEs), long tandem repeats (LTR), DNA transposons (DNA), unclassified (unknown) repeat families, small RNA repeats (SmRNA), and others (consisting of small, but classified repeat groups). The total is the total percentage of base pairs made up of repeats in each genome assembly, respectively.

| Assembly | LINE | SINE | LTR | DNA | Unclassified | SmRNA | Others | Total (%) |
| --- | --- | --- | --- | --- | --- | --- | --- | --- |
| *Candidula unifasciata* | 1,253,318 | 427,509 | 11,975 | 298,828 | 1,334,718 | 413,197 | 708,740 | 61.1 |
| *Cepaea nemoralis* | 2,820,864 | 342,120 | 209,476 | 443,363 | 4,400,828 | 444,489 | 1,267,814 | 77.0 |

Approximately one third of the assembled genome (33.96%) was identified as Transposable elements (TEs) such as long interspersed nuclear elements (LINEs; 25.03%), short interspersed nuclear elements (SINEs; 4.23%), long tandem repeats (LTR; 0.60%) and DNA transposons (4.10%).

We predicted 22,464 genes in the *C. unifasciata* genome (Table 3) by using a homology-based gene prediction and EVM. Among the identified proteins, 13,221 (62.27%) were annotated to have at least one GO term. Finally, 21,231 proteins (94.51%) were assigned to at least one of the database from InterProScan (Table 3). BUSCO and functional annotations results indicated high quality.

**Table 3.** Functional annotation of the predicted protein-coding genes for *C. unifasciata* genome.

|  |  | *C. unifasciata* |
| --- | --- | --- |
| **Number** |  |  |
|  | **Gene** | 22,464 |
|  | **mRNA** | 22,464 |
|  | **Exon** | 147,783 |
|  | **CDS** | 147,783 |
| **Mean** |  |  |
|  | **mRNAs/gene** | 1 |
|  | **CDSs/mRNA** | 6.58 |
| **Median length** |  |  |
|  | **Gene** | 11,931 |
|  | **mRNA** | 11,931 |
|  | **Exon** | 129 |
|  | **Intron** | 2,025 |
|  | **CDS** | 129 |
| **Total space (Mb)** |  |  |
|  | **Gene** | 379,573,459 |
|  | **CDS** | 26,582,739 |
| **Single** |  |  |
|  | **CDS mRNA** | 3,562 |
| **InterproScan** |  | 21,231 (94.51%) |
| **GO** |  | 13,221 (62.27%) |
| **Reactome** |  | 5,069 (22.56%) |
| **SwissProt** |  | 16,809 (74.83%) |

**Table 4.**
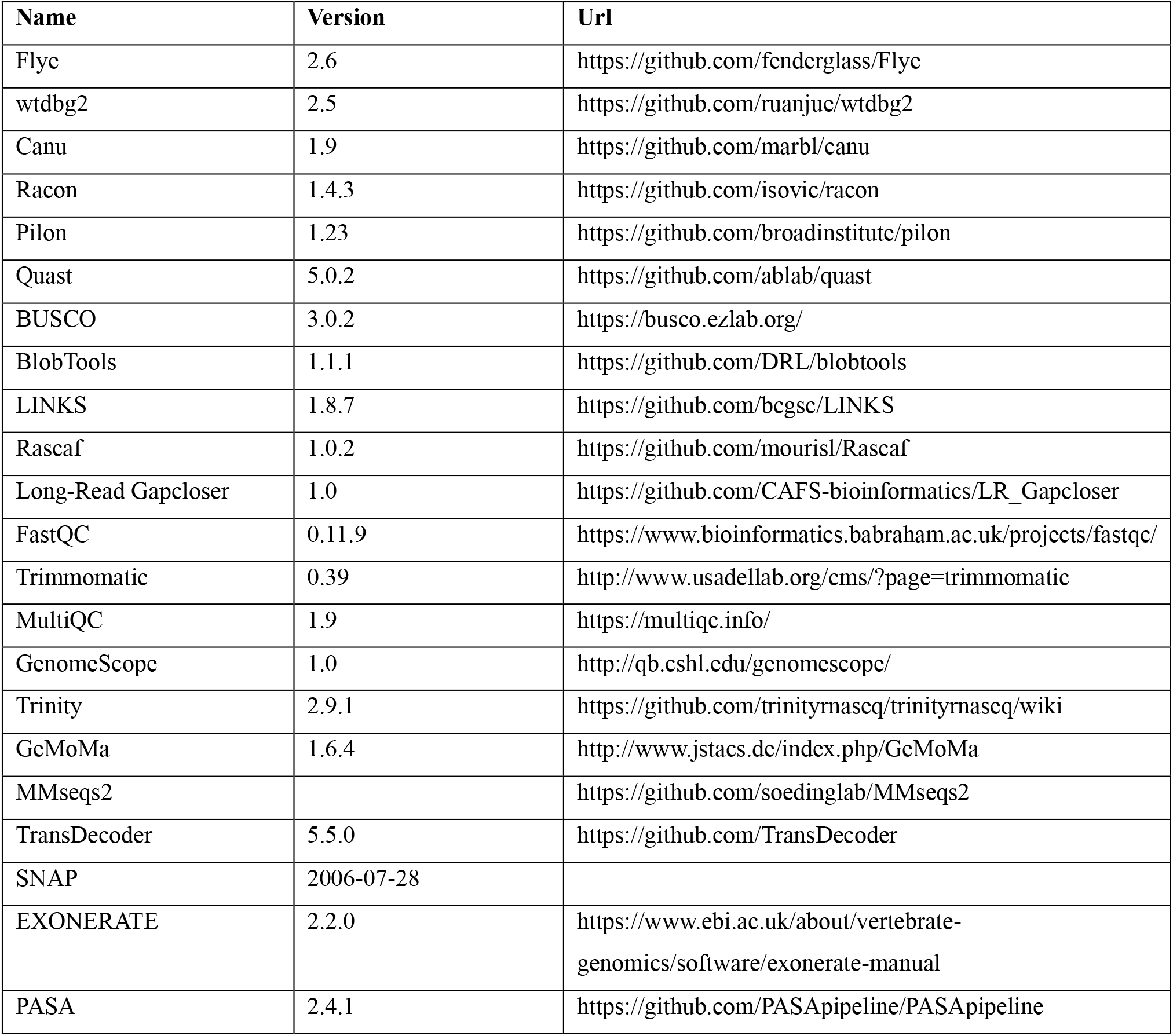
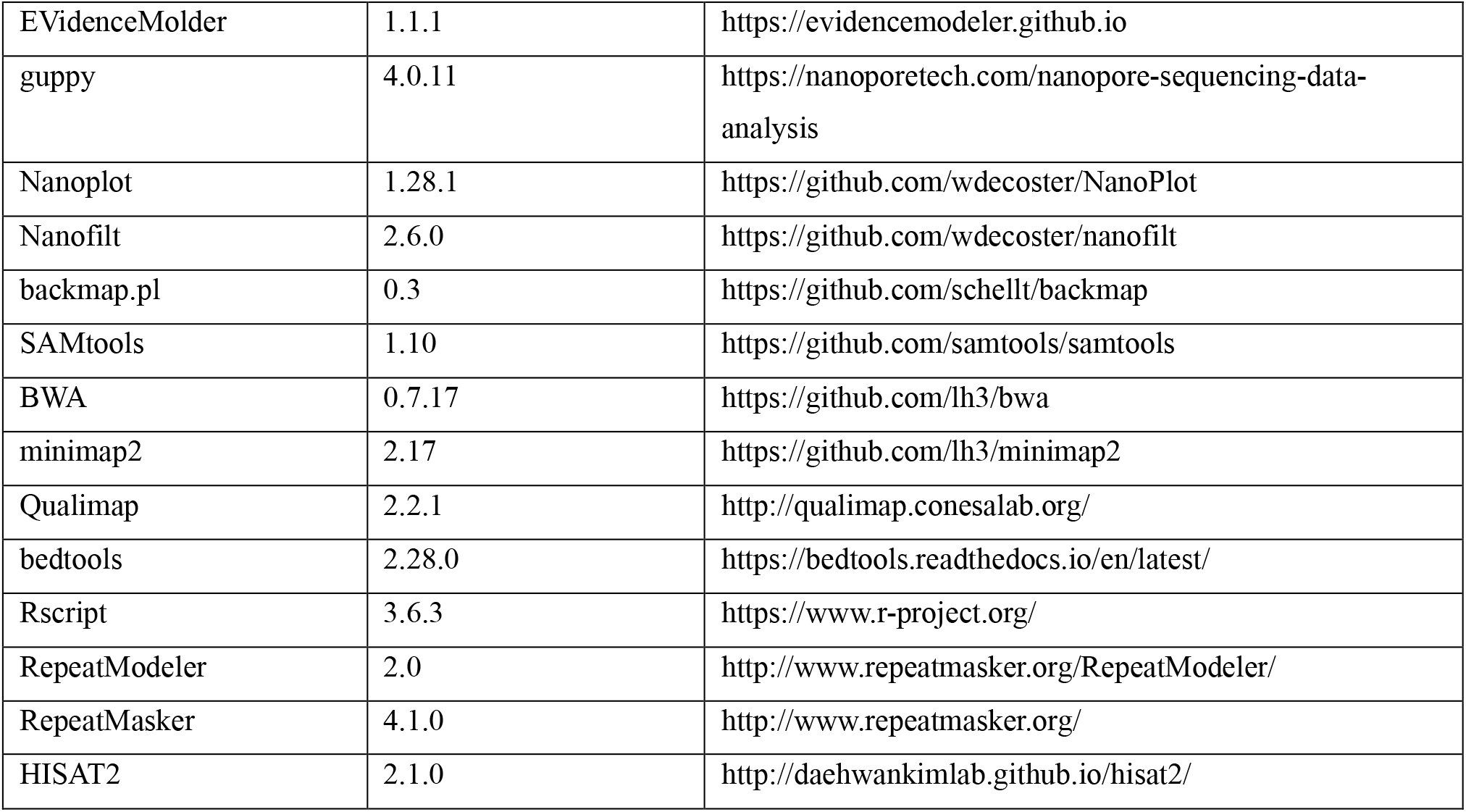
Software employed in this work, their package version and source availability.

Total protein-coding genes was in the range of other gastropods annotations (Schell *et al*. 2017; Liu *et al*. 2018; Guo *et al*. 2019), however this number represented only the half of its closest relative *Cepaea nemoralis* (Saenko *et al*. 2021).

## 4. Conclusions

Here, we present a draft assembled and annotated genome of the land snail *Candidula unifasciata*. The obtained genome is comparable with other land snail and Gastropoda genomes publicly available. The new genome resource will be reference for further population genetics, evolutionary and genomic studies of this highly world-wide diverse group.

## Data Availability Statement

All raw data generated for this study (Illumina, PacBio, MinION, and RNA-seq reads) are available at the European Nucleotide Archive database (ENA) under the Project number: PRJEB41346. The final genome assembly and annotation can be found under the accession number GCA_905116865.

## Acknowledgments

This work was funded by LOEWE-Centre for Translational Biodiversity Genomics (LOEWE-TBG). We thank Damian Baranski for help with the DNA isolation and library preparations. Luis J. Chueca was supported by a Post-doctoral Fellowship awarded by the Department of Education, Universities and Research of the Basque Government (Ref.: POS_2018_1_0012).

## Author contributions

M.P. and L.J.C. conceived the idea. M.P. collected the specimens. L.J.C. designed and performed the bioinformatic analyses with support of T.S. L.J.C. prepared the manuscript, and all authors edited and approved the final version.

## Figures and Tables

**Table S1.**
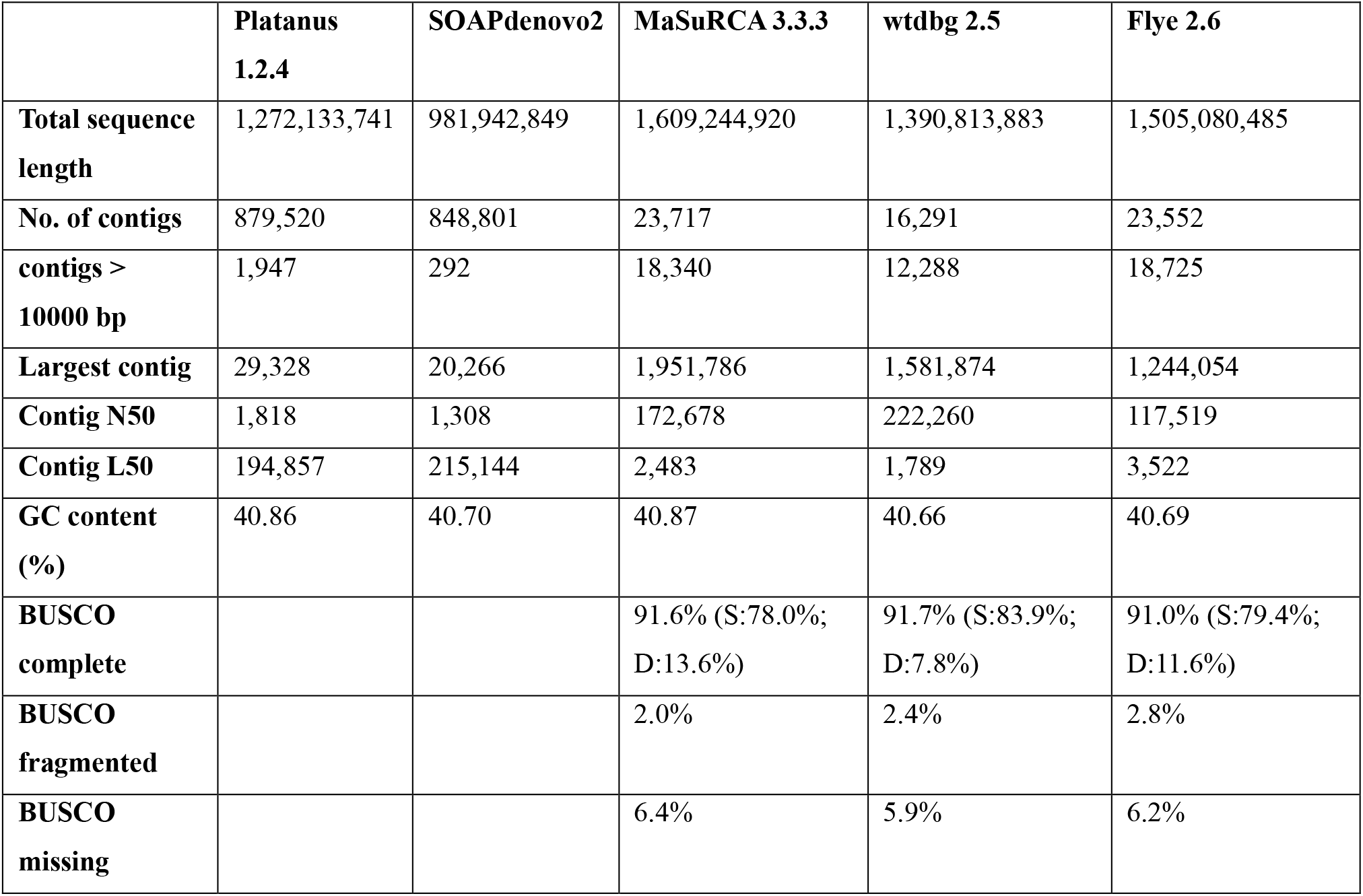
Comparison between draft genomes assemblies obtained by the different tools.

|  | **Platanus**<br>**1.2.4** | **SOAPdenovo2** | **MaSuRCA 3.3.3** | **wtdbg 2.5** | **Flye 2.6** |
| --- | --- | --- | --- | --- | --- |
| **Total sequence length** | 1,272,133,741 | 981,942,849 | 1,609,244,920 | 1,390,813,883 | 1,505,080,485 |
| **No. of contigs** | 879,520 | 848,801 | 23,717 | 16,291 | 23,552 |
| **contigs > 10000 bp** | 1,947 | 292 | 18,340 | 12,288 | 18,725 |
| **Largest contig** | 29,328 | 20,266 | 1,951,786 | 1,581,874 | 1,244,054 |
| **Contig N50** | 1,818 | 1,308 | 172,678 | 222,260 | 117,519 |
| **Contig L50** | 194,857 | 215,144 | 2,483 | 1,789 | 3,522 |
| **GC content (%)** | 40.86 | 40.70 | 40.87 | 40.66 | 40.69 |
| **BUSCO complete** |  |  | 91.6% (S:78.0%; D:13.6%) | 91.7% (S:83.9%; D:7.8%) | 91.0% (S:79.4%; D:11.6%) |
| **BUSCO fragmented** |  |  | 2.0% | 2.4% | 2.8% |
| **BUSCO missing** |  |  | 6.4% | 5.9% | 6.2% |

